# Modulated efficacy CRISPRi reveals evolutionary conservation of essential gene expression-fitness relationships in bacteria

**DOI:** 10.1101/805333

**Authors:** John S. Hawkins, Melanie R. Silvis, Byoung-Mo Koo, Jason M. Peters, Marco Jost, Cameron C. Hearne, Jonathan S. Weissman, Horia Todor, Carol A. Gross

## Abstract

Essential genes are the central hubs of cellular networks. Despite their importance, the lack of high-throughput methods for titrating their expression has limited our understanding of the fitness landscapes against which essential gene expression levels are optimized. We developed a modified CRISPRi system leveraging the predictable reduction in efficacy of imperfectly matched sgRNAs to generate specific levels of CRISPRi activity and demonstrate its broad applicability in bacteria. Using libraries of mismatched sgRNAs, we characterized the expression-fitness relationships of essential genes in *Escherichia coli* and *Bacillus subtilis*. Remarkably, these relationships co-vary by pathway and are predominantly conserved between *E. coli* and *B. subtilis* despite ~ 2 billion years of evolutionary separation, suggesting that deeply conserved tradeoffs underlie bacterial homeostasis.

**One Sentence Summary:** Bacterial essential genes have varying responses to CRISPRi knockdown that are largely conserved across ~2 billion years of evolution.

## Main Text

Bacteria must optimize protein production to maximize survival and growth in constantly changing environments. Given the high energetic cost of protein synthesis, optimizing expression is particularly important for essential genes: although only ~5-10% of the genome, they constitute a disproportionate fraction (~50%) of the proteome (*1*) and insufficient expression is, by definition, fatal. Previous work using promoter replacement revealed gene-, environment-, and antibiotic-specific fitness effects of altering essential gene expression (*2*–*7*), but the lack of a facile method for systematically perturbing bacterial gene expression has thus far prevented a comprehensive understanding of how bacteria optimize expression of their essential protein complement. CRISPR interference (CRISPRi), which blocks bacterial transcription by targeting a catalytically dead Cas9 (dCas9) to a gene using a complementary sgRNA, has been used to perturb essential gene expression in its native context. However, tuning transcriptional repression by adjusting dCas9 or sgRNA abundance (*8*, *9*) is noisy and precludes the interrogation of multiple knockdown levels in a single experiment (*10*). Building on previous work (*10*, *11*), we reasoned that we could instead modulate transcriptional repression by programming a highly expressed CRISPRi system with sgRNAs imperfectly matched to their target. This would allow us to explore the fitness landscape of essential gene expression by enabling massively parallel interrogation of the fitness effects of multiple levels of CRISPRi activity across genes in a single pooled growth experiment.

We first explored how mismatches affect sgRNA activity by generating a comprehensive library of sgRNA spacers targeting *gfp* (3201 total), consisting of all spacers fully complementary to the non-template strand (33), a majority of their possible single mismatch variants (47/60), and a subset of their possible double mismatch variants (49/1710) (Fig. S1A). Using FACS-seq (Fig 1A, Methods), we quantified the ability of these sgRNAs to repress transcription of a highly expressed chromosomal copy of *gfp* both in *E. coli* and *B. subtilis* (Fig S2A-C and table S1). We found that sgRNAs with either single (Fig. 1B) or double (Fig. S3A) mismatches in their base-pairing regions generated the full range of repression (no efficacy to full efficacy) in both species. Importantly, sgRNA activity was unimodal (Fig. S2E-H) and highly correlated between *E. coli* and *B. subtilis* (R^2^: singly mismatched sgRNAs = 0.65, doubly mismatched sgRNAs = 0.61, all sgRNAs = 0.71, Fig. 1B, fig. S3A, and table S1), despite an evolutionary distance of several billion years and differences in experimental setup (*E. coli:* plasmid-encoded sgRNAs, *B. subtilis:* chromosomally integrated sgRNAs). This suggests that the primary determinant of CRISPRi efficacy in bacteria is the interaction between the dCas9-sgRNA complex and DNA, rather than organism-specific factors such as the host’s transcriptional machinery.

**Fig. 1.**
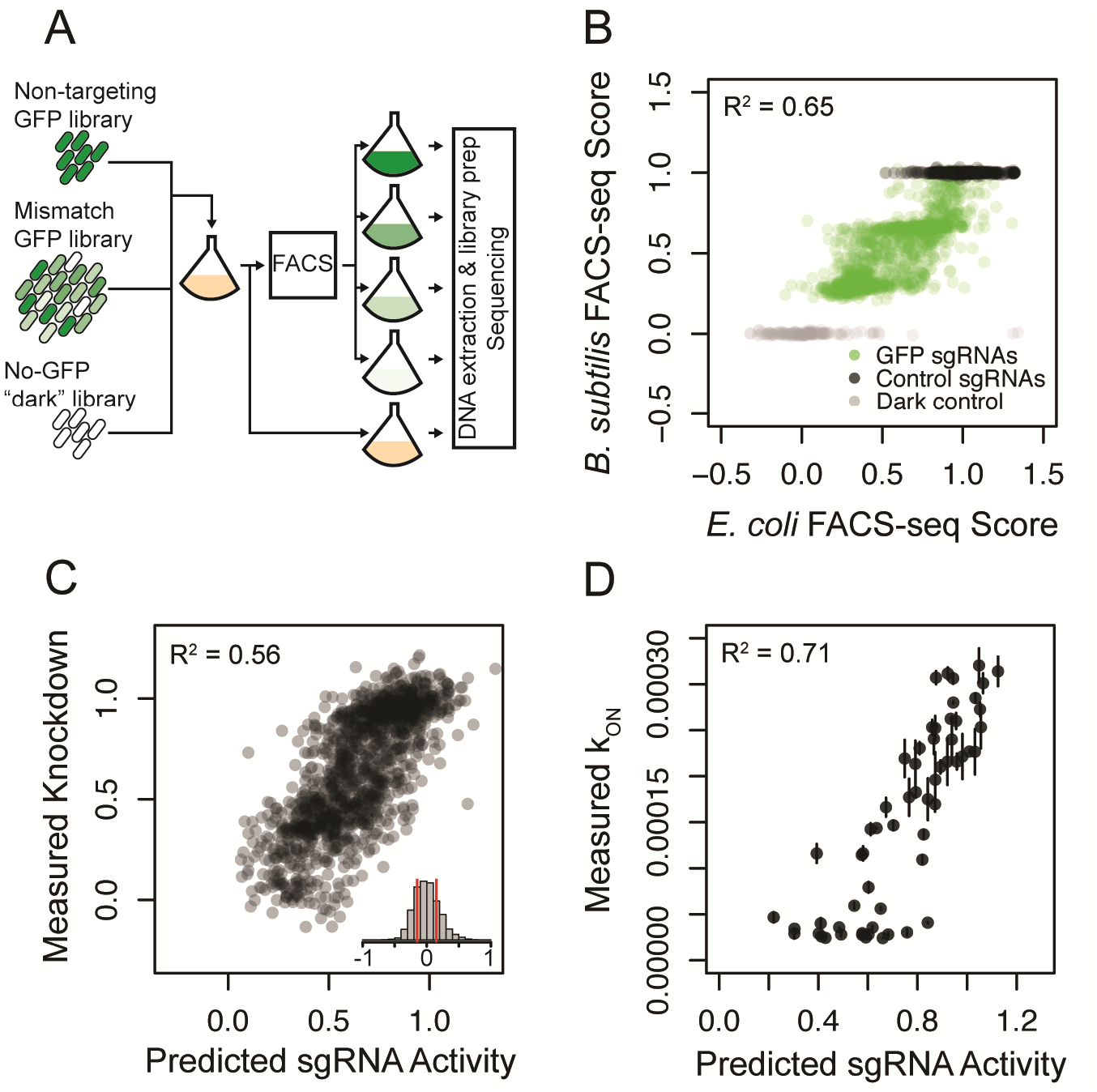
Singly mismatched sgRNAs reproducibly generate a range of knockdown efficacies in *B. subtilis* and *E. coli* and are accurately predicted by a simple linear model. (A) Workflow of a FACS-seq experiment. (B) FACS-seq scores (average of 2 biological replicates) for each singly mismatched sgRNA targeting *gfp* in *B. subtilis* and *E. coli*. Additional noise in *E. coli* likely represents changes in plasmid copy number during outgrowth. (C) The predictions of a linear model trained on GC%, mismatch position, and mismatch identity compared to the measured relative *gfp* knockdown efficacies of each sgRNA averaged over both species. Inset is a histogram of the differences between predicted and measured knockdown, reflecting both prediction and measurement error: 56% of sgRNAs measured within 0.15 of their predicted activity (red bars). (D) The predictions of the linear model compared to the measured singly mismatched sgRNA association rates (k_ON_) in vitro (*14*).

Given the species-independent performance of *gfp-*targeting mismatched sgRNAs, we next asked whether we could accurately predict the effects of single mismatches on sgRNA activity. Informed by previous work on CRISPRi off-target effects (*11*, *12*) and concurrent work on mismatched sgRNAs in a mammalian context (*13*), we constructed a linear model using the position and base substitution of the mismatch and the GC% of the fully complementary spacer as features. We trained this model on the *E. coli, B. subtilis* (Fig. S4 and fig. S5), or species-averaged relative efficacy of our *gfp*-targeting singly mismatched sgRNAs (Fig. 1C) and found that the effects of single mismatches could be robustly predicted in all cases (species-averaged R^2^ = 0.56, 11-fold CV-MSE = 0.10 ^+^/_-_ 0.08). Assuming that mismatches have independent effects on sgRNA efficacy, the model also accurately predicted double mismatch efficacy (R^2^ = 0.53, Fig. S3B). To further validate this model, we compared our predicted sgRNA activity to the previously measured association rates (k_on_) of a dCas9-sgRNA complex to 60 singly mismatched and 1130 doubly mismatched DNA sequences (*14*) (Fig. 1D and fig. S3C). Our predicted efficacy was highly correlated (R^2^: single mismatches = 0.71, double mismatches = 0.45) to the k_on_ measured in this *in vitro* system, supporting the hypothesis that mismatched-CRISPRi functions by reducing the association rate of the dCas9-sgRNA complex for the target DNA. Consistent with this idea, our model recapitulates many biophysical properties of RNA-DNA interactions such as the relative stability of rG:dT basepairs implied by its high coefficient for A to G transitions (*15*) (Table S2 and fig. S2D). Taken together, these data strongly suggest that a simple linear model trained on the relative efficacy of our *gfp*-targeting singly mismatched sgRNA library can be used to design mismatched sgRNAs with a specific activity level targeting any gene.

Using our model of mismatched sgRNA activity, we designed a set of sgRNAs targeted to the essential gene complement of *E. coli* and *B. subtilis* (~300 genes in each species, Table 3) and predicted to have a range of activity. We generated large pooled libraries of strains in which each essential gene is targeted by 100 sgRNAs (10 fully matched guides each with 9 singly mismatched variants, Methods, Fig. S1C) and compact libraries in which each essential gene is targeted by 11 sgRNAs (*44*). Additionally, for two well characterized essential genes encoding UDP-GlcNAc-1 carboxyvinyltransferase (*E. coli: murA*, *B. subtilis: murAA*), and dihydrofolate reductase (*E. coli: folA*, *B. subtilis: dfrA*), we generated comprehensive libraries (at least 47/60 single mismatch variants for each sgRNA within the gene, Methods, Fig. S1B). The libraries were grown for 10 doublings, maintaining exponential phase through back-dilution (Fig. 2A). We calculated the relative fitness (*16*, *17*) of each strain by comparing its relative abundance (quantified by next-generation sequencing of the sgRNA spacers) to the relative abundance of 1000 non-targeting sgRNAs at the start and end of each experiment (Methods, Table S3). Relative fitness is defined as the fraction of doublings a strain undergoes compared to wild-type over the course of the experiment. Strains with a relative fitness of 1 grow as well as wild-type; lower values imply slower growth. Relative fitness was highly reproducible in both species (R^2^ > 0.9, Fig. S6A-B), was validated by orthogonal measurements of individual strain fitness (Fig. S6C), and was consistent within fully complementary sgRNAs targeting the same gene (Sup. Text 1). Our relative fitness values were correlated with previously reported measurements (*18*,*19*) but had greatly expanded dynamic range due to greater sequencing depth and a shorter growth period (Fig. S6D-E, Sup. Text 2). This expanded dynamic range enabled measurement of negative relative fitness, which indicates active depletion from the pool. CRISPRi targeting of 23 *E. coli* genes and 24 *B. subtilis* genes reproducibly (>5 sgRNAs) caused negative relative fitness (Table S4). Consistent with an interpretation of negative relative fitness as lysis, a majority (15/24) of these *B. subtilis* genes caused lysis when targeted with a fully complementary sgRNA (*8*) (Table S4, Methods).

**Fig. 2.**
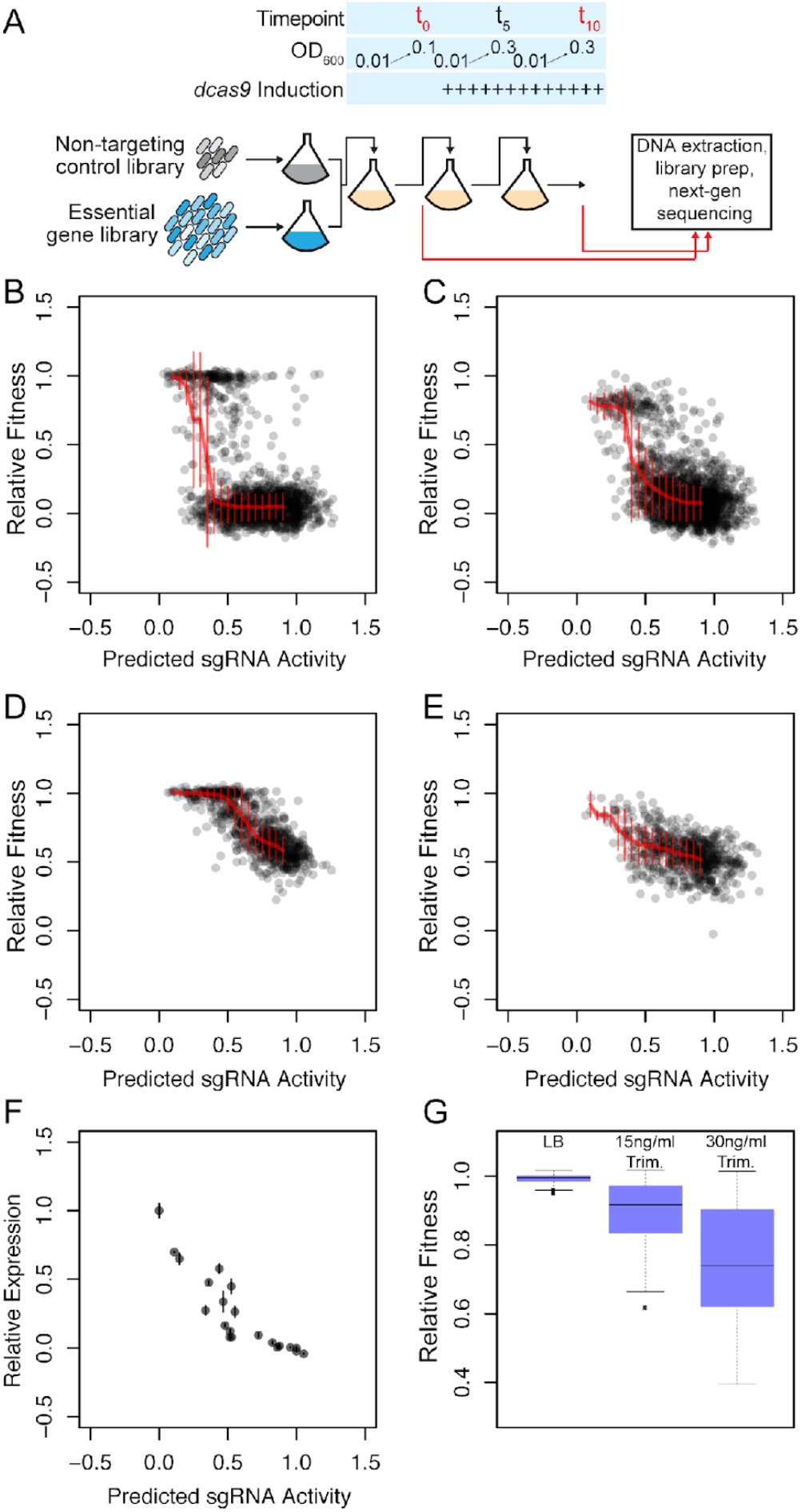
Singly mismatched sgRNAs targeting two essential genes in *E. coli* and *B. subtilis* illustrate the gene-specific nature of expression-fitness relationships. (A) Schematic of the fitness experiment design. (B-E) Predicted sgRNA activity and measured relative fitness of singly mismatched sgRNA targeting (B) *murAA* in *B. subtilis*. (C) *murA* in *E. coli*. (D) *dfrA* in *B. subtilis*. (E) *folA* in *E. coli*. Median and SD values of relative fitness for sgRNAs grouped into 17 bins based on predicted knockdown are shown in red. (F) Predicted sgRNA activity and relative expression for 18 singly mismatched sgRNA targeting a *murAA-gfp* transcriptional fusion in a *murAA*–complemented *B. subtilis* strain (Methods). Relative expression is shown as the median single-cell GFP fluorescence, normalized as a fraction of control (no sgRNA). (G) Measured relative fitness of singly mismatched sgRNAs targeting *dfrA* in *B. subtilis* with relative fitness > 0.95 in LB (n=252). Measured fitness values of the same sgRNAs in the presence of sub-inhibitory concentrations of trimethoprim.

We next assessed whether comparing the predicted activity of sgRNAs to their relative fitness would allow us to infer per gene expression-fitness relationships. First, we asked whether predicted sgRNA activity was inversely correlated to relative fitness across our data set. As expected, we found strong negative correlations both within sgRNA families (sgRNAs targeting the same locus) and within genes (Fig. S7, Methods). Weaker correlations in *E. coli* likely reflect variation in sgRNA plasmid copy number and/or *E. coli* specific effects (*20*). Second, we examined the expression-fitness relationships of *murA/murAA* and *folA/dfrA* using comprehensive mismatched sgRNA libraries. Consistent with previous studies (22-24), we find that CRISPRi targeting of *murA/murAA* bimodally affects fitness (Fig. 2B-C), while CRISPRi targeting of *folA/dfrA* linearly affects growth rate above an initial threshold of activity (Fig. 2D-E). Highlighting the approximately linear effect of *folA/dfrA* repression on fitness, our model could accurately predict *gfp* knockdown after being trained on the fitness effects of mismatched sgRNAs targeting those genes (Fig. S4 and fig. S5). Third, we measured the ability of 18 mismatched sgRNAs to repress a *murAA-gfp* transcriptional fusion in *B. subtilis*. To enable quantification of lethal levels of knockdown and to minimize transcriptional feedback these measurements were conducted in a *B. subtilis* strain complemented with non-targeted *murAA* (Methods). The predicted activity of sgRNAs targeting *murAA* in this experiment closely tracked actual knockdown (Fig. 2F), suggesting that the non-linear expression-fitness relationship of *murAA* without complementation (Fig. 2B) reflect non-linearly decreasing growth due to MurAA depletion, transcriptional feedback, cell lysis, or other host specific effects. Finally, we confirmed that low efficacy (relative fitness > 0.95) sgRNAs targeting *dfrA* were functional by measuring the fitness of the *B. subtilis dfrA* library in the presence of trimethoprim, a direct inhibitor of DfrA. Trimethoprim decreased the fitness of low efficacy sgRNAs targeting *dfrA* suggesting that these sgRNAs impact *dfrA* expression even in the absence of a measurable fitness defect (Fig. 2G). Taken together, these validation experiments strongly suggest that we can accurately and sensitively probe the expression-fitness relationships of essential genes in *E. coli* and *B. subtilis* by comparing the predicted activity of mismatched sgRNAs to their measured fitness using a pooled screening approach.

Examining the essential gene expression-fitness relationships, we were struck by their diverse and gene-specific nature (Fig. S8, fig. S9, and table S3). To quantitatively characterize these differences, we first binned the sgRNAs targeting each gene according to predicted sgRNA activity and calculated the median fitness within each bin (Methods, Fig. 2B-E, table S5). Next, we used these simplified representations of per gene expression-fitness relationships to calculate pairwise distances between *E. coli* and *B. subtilis* essential genes. Within each organism, we found that the expression-fitness relationships of genes involved in the same biological process (whether defined by KEGG, GO biological process, or COG) were significantly more similar to each other than to those of genes involved in different biological processes, even when excluding gene pairs in the same operon to account for CRISPRi polarity (all *p* < 10^−16^, Methods). Inversely, clustering genes by the shape of their expression-fitness curves produced functional enrichments (Table S6) in both *E. coli* and *B. subtilis*. Finally, in a cross-species comparison, the expression-fitness curves of essential genes were, as a group, more similar (*p* < 10^−10^) to that of their homologs than to other genes in the opposing species. Taken together, these data suggest that these expression-fitness curves are both biologically meaningful and representative of deeply conserved homeostatic constraints on bacterial physiology.

To explore the conserved optimizations of bacterial essential gene expression, we examined three functional categories having similar expression-fitness relationships in *E. coli* and *B. subtilis*. CRISPRi targeting of essential cofactor biosynthesis genes (KEGG pathways under “Metabolism of cofactors and vitamins”) did not strongly affect fitness in either species after 10 generations (Fig. 3A-B). This observation is consistent with the small-colony but non-culturable phenotype of essential cofactor biosynthesis gene deletions (*21*) and suggests that these cofactors and/or the enzymes producing them are present in excess of what is required for exponential growth. This buffer may be required to enable rapid shifts in metabolism in response to changing environmental conditions, similar to what has been proposed for the pentose-phosphate pathway (*22*).

**Fig. 3.**
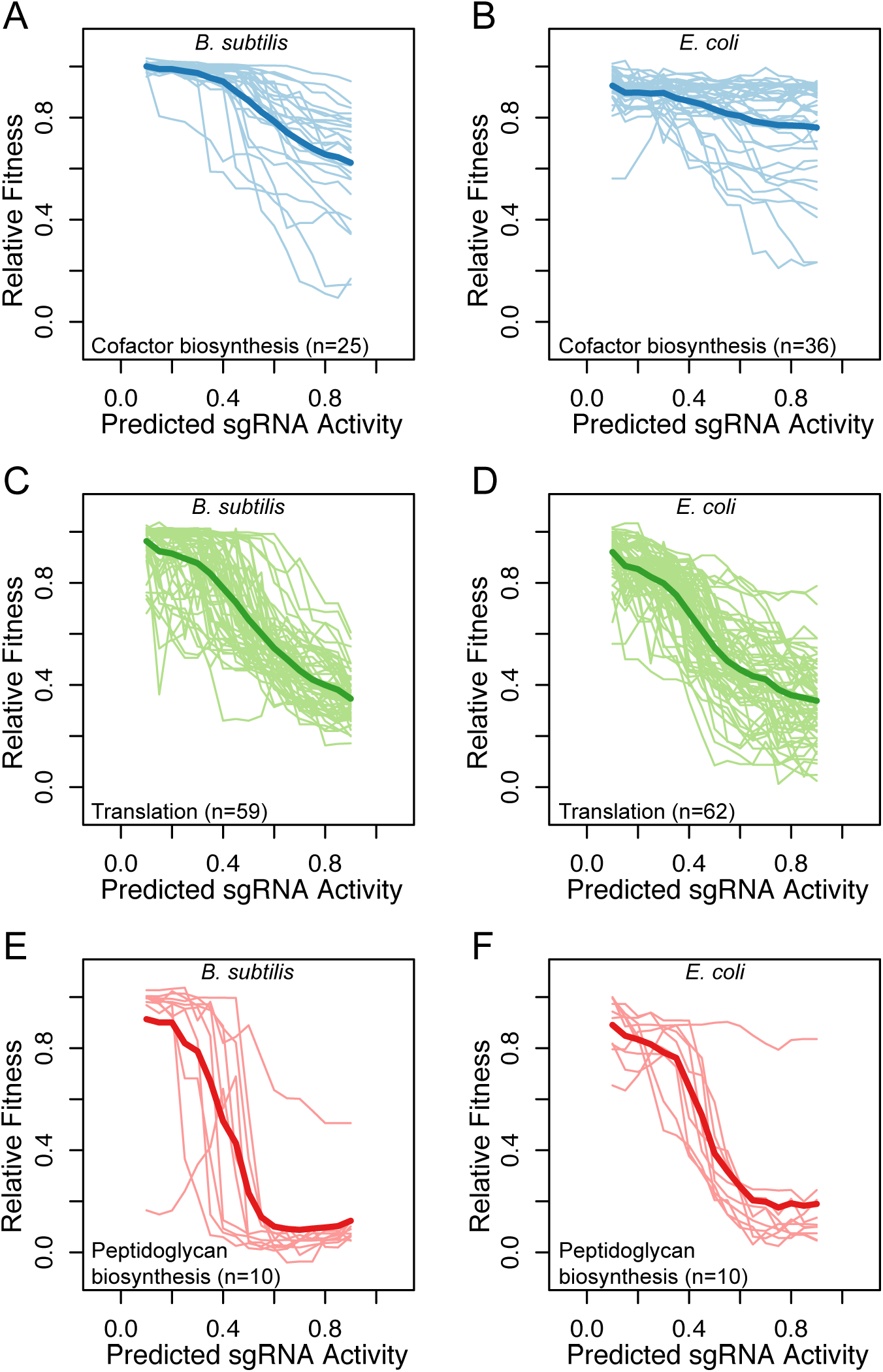
Expression-fitness relationships of essential genes are conserved within biological process and between *B. subtilis and E. coli.* Relative fitness compared to predicted knockdown for: essential cofactor biosynthesis genes (KEGG pathways under “Metabolism of cofactors and vitamins”) in *B. subtilis* (A) or *E. coli* (B); KEGG pathways under “Translation” in *B. subtilis*(C) or *E. coli* (D); peptidoglycan biosynthesis (KEGG pathway ko00550) in *B. subtilis*. (E) or *E. coli* (F).

The robustness of both bacteria to CRISPRi targeting of essential cofactor synthesis genes contrasts with the strong, approximately linear effect of targeting genes involved in translation (KEGG pathways under “Translation”, Fig. 3C-D). Previous work has established a linear relationship between growth rate and the number of ribosomes per cell during exponential growth in *E. coli, B. subtilis*, and other bacteria (*23*–*25*). By linearly inhibiting ribosomal protein expression, we likely decrease the number of functional ribosomes, leading to a corresponding linear decrease in growth rate. Moreover, feedback to restore ribosomal protein expression is unlikely because most ribosomal proteins are negatively regulated by their excess relative to rRNA (*26*, *27*). Depletion of translation factors has a similarly linear effect on growth rate (Fig. S8 and fig. S9), likely due to slowed elongation rate (*28*) as has been shown for some translation inhibitors (*24*). The conserved linear relationship between the expression of proteins involved in translation and growth rate reinforces the universal importance of translational capacity for determining growth rate.

CRISPRi targeting of genes involved in cytoplasmic peptidoglycan precursor synthesis (KEGG ko00550) also generated strong phenotypes in both species. However, in contrast to the linear expression-fitness relationship of genes involved in translation, peptidoglycan synthesis genes exhibited bimodal fitness outcomes that depended on predicted sgRNA activity (Fig. 3E-F). Cells tolerated partial repression of these genes without exhibiting a fitness defect, perhaps due to transcriptional feedback and/or an excess of enzyme. If expression was sufficiently repressed, these strains lysed (Table S4) as has been described for *murA, murG*, and *mraY* inhibition in *E. coli (29–31)* and for *murC, murD*, and *murG* depletion in *B. subtilis (8)*. However, the dearth of intermediate fitness outcomes upon repression of peptidoglycan precursor synthesis in both species is surprising. It suggests that neither species is able to slow growth rate in response to reduced flux through cytoplasmic peptidoglycan precursor synthesis to prevent lysis. It has been proposed that bacteria use peptidoglycan precursor concentration to sense and balance cellular metabolism and growth (*32*). This would be incompatible with direct feedback regulation of cytoplasmic peptidoglycan precursor synthesis and may explain the sharp transition between growth and lysis.

Given the similarity between the expression-fitness curves of most essential genes in *E. coli* and *B. subtilis*, we reasoned that homologs with substantially different expression-fitness curves may illustrate biologically relevant differences between the two organisms. We identified 9 homologs as significantly different between the two organisms (Table S7, FDR < 0.2), most of which encoded enzymes involved in peptidoglycan synthesis and maturation. In contrast to the conserved bimodal expression-fitness relationships of genes involved in cytoplasmic peptidoglycan precursor synthesis just discussed (Fig. 4, group 3), CRISPRi targeting of genes required for producing either UDP-GlcNAc (Fig. 4, group 1) or *meso-*DAP (Fig. 4, group 2) differentially affected fitness in the two species*. E. coli* was robust to CRISPRi targeting of these genes, while *B. subtilis* was sensitive, lysing when these genes were targeted with high activity sgRNAs (Fig. 4, table S4, and table S7). The differential effect of CRISPRi targeting on these genes could be attributed to buffering of either expression or activity in *E. coli* but not in *B. subtilis*, perhaps mediated by divergent regulatory mechanisms (*33*, *34*). Differences in the fitness effect of DAP pathway knockdowns may also be accounted for by peptidoglycan stem-peptide recycling through the *E. coli* enzyme Mpl (*35*). *E. coli* was also significantly more tolerant of perturbation of *mreBCD* than *B. subtilis* (Fig. 4, group 4), exhibiting a minimal fitness defect after 10 generations (but lysis after 15 generations, Table S3). This observation is consistent with the small effect of CRISPRi targeting of *mrdA* (the PBP2 associated with MreBCD) on fitness in *E. coli* (Fig. S4), and with previous work which found that *Enterobacter cloacae* is also relatively unaffected by *mreBCD* CRISPRi targeting (*36*). It is unclear why *E. coli* and other Gram-negative bacteria are less affected by *mreBCD* CRISPRi targeting than *B. subtilis*, however transcriptional buffering through feedback may play a role. Alternatively, the substantially higher turgor pressure in *B. subtilis (37)* may make it less tolerant of cell wall abnormalities.

**Fig. 4.**
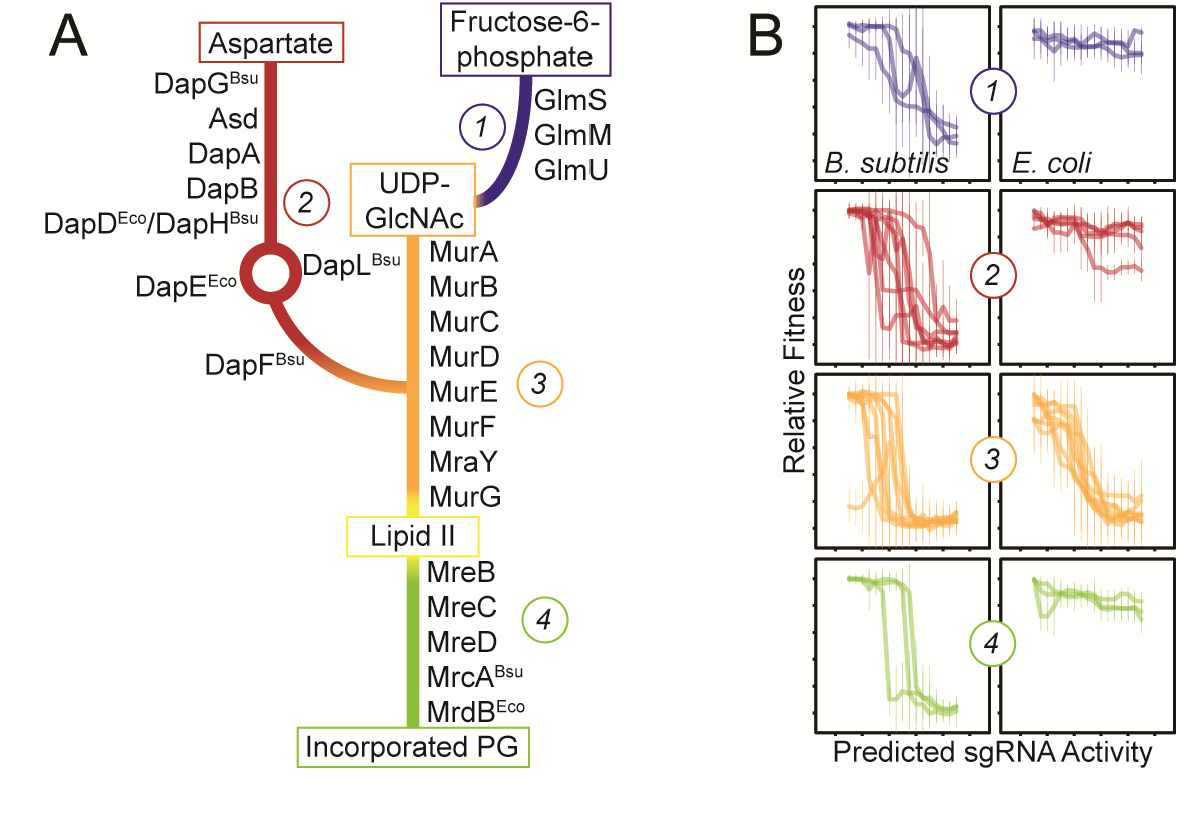
Similar and different expression-fitness relationships of cell wall biosynthesis genes in *B. subtilis* and *E. coli*. (A) Pathway of peptidoglycan synthesis and incorporation, color coded by portion of the pathway. (B) Predicted knockdown vs. relative fitness for the groups of essential genes from pathway sections indicated in (A), in *B. subtilis* and *E. coli*.

Singly mismatched CRISPRi is a universal approach for systematically perturbing bacterial gene expression. Leveraging this technique, we explore the expression-fitness relationships of essential genes in *E. coli* and *B. subtilis* and reveal that the basic biological constraints driving essential gene fitness landscapes are conserved over >2 billion years of evolution. These studies inform target selection for drug design, illuminate aspects of bacterial growth, and provide a starting point for investigating how bacteria program robustness into their essential gene network.

## Acknowledgements

We thank M. Kampmann, D. Santos, M. Horlbeck, A. Banta, G-W Li, and members of the J.S.W. and C.A.G. Labs for extensive helpful discussions, C. Lu, C. Liem, M. DeVera, and R. Pak for assistance with library cloning, J. Garabino for help with flow cytometry, and E. Chow, D. Bogdanoff, and K. Chaung from the UCSF Center for Advanced Technology for help with sequencing.

## Author contributions

JSH, MRS, BMK, JMP, HT and CAG designed the study; JSH, MRS, BMK, and CCH performed experiments; JSH, MRS, HT, MJ built software; JSH, MRS, BMK, HT analyzed data; JSH, MRS, BMK, JMP, HT, MJ, JSW and CAG wrote and/or edited the manuscript.

## Data and materials availability

All sequencing data have been deposited in the Short Read Archive with BioProject accession PRJNA574461. All analyzed data is available in the main text or the supplementary materials. Unpublished code used in this manuscript is available on GitHub (https://github.com/traeki/mismatch_crispri and https://github.com/traeki/sgrna_design).

## Supplementary Materials

Materials and Methods

Figures S1-S11

Tables S1-S10

References (38-43)

