## Supplementary Materials for "Modulated efficacy CRISPRi reveals evolutionary conservation of essential gene expression-fitness relationships in bacteria"

**This PDF file includes:**

Materials and Methods
Supplementary Text
Figs. S1 to S11
Tables S1 to S10

### Materials and Methods

Materials

*Bacillus subtilis* strain construction and growth conditions

All primers are described in Table S8, and all strains are described in Table S9.

All *B. subtilis* strains were constructed in the wildtype 168 background using natural competence as previously described (*21*). For all individual CRISPRi strains and libraries, a recipient strain encoding *dcas9* under control of the P*xyl* promoter at the *lacA* locus (strain CAG74209) (*8*), was transformed with an sgRNA plasmid (see "sgRNA plasmid construction") which recombines in single copy at the *amyE* locus, selecting for chloramphenicol resistance. In select cases, single- vs. double-crossover events from plasmid integration were distinguished by streaking on starch plates to assay disruption of *amyE*.

For the GFP knockdown FACS-seq experiments, two modified recipient strains expressing *dcas9* were constructed: one encoding *gfp* (strain CAG78920) and the other encoding *rfp* (strain CAG78921). To construct these, the *dcas9* strain (strain CAG74209) was transformed with pDG1731-gfp or pDG1731-rfp to integrate P*veg-gfp-spc* or P*veg-rfp-spc*, respectively, at the *thrC* locus, selecting for spectinomycin resistance. All subsequent transformations of the *gfp* and *rfp*-marked strains required threonine supplementation in the competence media (40μg/ml), as *thrC* is disrupted.

For flow cytometry-based competition experiments (see "Relative fitness validation"), the *dcas9* recipient strain was transformed with a modified sgRNA plasmid that also encodes either P*veg-gfp* or P*veg-rfp* (see "sgRNA plasmid construction").

A *murAA-gfp* transcriptional fusion knock-down reporter strain was constructed by transformation of the *dcas9* strain (above) with the DNA fragments containing constitutively expressed *rfp* with removable $kan^R$ cassette, *murAA-gfp* transcriptional fusion with removable $kan^R$ cassette, and constitutively expressed non-targeted *murAA* with spectinomycin resistant gene, of which fragments were integrated into *sacA*, *murAA* and *thrC* locus respectively, sequentially in that order. The *B. subtilis* P*veg* promoter was used for constitutive expression of *rfp* and non-targeted *murAA*. The DNA fragment containing constitutively expressed *rfp* with removable $kan^R$ cassette was constructed by joining PCR of three fragments: P*veg-rfp-kan*$^R$ fragment amplified from pACYC-*rfp-kanR*, and 1kb each 5′ and 3′ flanking sequences of *sacA*. The DNA fragment containing *murAA-gfp* transcriptional fusion with removable $kan^R$ cassette was constructed by the joining of *gfp-kanR* amplified from pACYC-gfp-kanR, 1kb of the 3' end of the *murAA* open reading frame, and 1kb downstream of the *murAA* open reading frame. Before the *gfp-kanR* fragment was integrated downstream of *murAA* to generate strain CAG78923, the *kanR* cassette was removed from *rfp* strain as described previously to generate strain CAG78922 (*21*). Non-targeted *murAA* was designed to remove PAM sequence or alter the sgRNA targeting sequence without substituting amino acid sequence of *murAA*. Non-targeted *murAA* DNA was generated by overlapping PCR with mutagenic primers (Table S8) and its *BsiWI/NruI* digested fragment were cloned into pJMP3 (Addgene #79875) digested with *BsrGI/PmeI*. The cloned plasmid was transformed into the *dcas9, rfp*, *murAA-gfp* strain, selecting for spectinomycin resistance, to generate strain CAG78924. Finally, this strain was transformed with sgRNA plasmids as described above.

Unless otherwise noted, all strain construction and growth assays for *B. subtilis* were done in LB medium and using antibiotics at the specified concentrations: erythromycin (1µg/ml), spectinomycin (100µg/ml), chloramphenicol (7.5µg/ml), kanamycin (7.5µg/ml).

*Escherichia coli* strain construction and growth conditions

All CRISPRi library strains were constructed in the wildtype BW25113 background by electroporating an sgRNA plasmid or plasmid pool (see "sgRNA plasmid construction") into a recipient strain encoding *dcas9* (for essential gene knockdown libraries), or *dcas9* and *gfp* or *rfp* (for GFP knockdown libraries), selecting for ampicillin resistance.

For the essential gene knockdown library recipient strain (strain CAG78830), Tn7 transposition was used to integrate a *dcas9* expression cassette into the Tn7att site using triparental mating of DAP(diaminopimelic acid)-dependent donors and selecting for gentamicin resistance in the absence of DAP, as previously described (*36*). The *dcas9* expression cassette is modified from previously described versions (*36*), contains *dcas9* from *S. pyogenes* (*12*)with a 3X Myc C-terminal tag, and is expressed from the IPTG-inducible promoter P*lLac-O1* (*38*) and regulated by *lacIq*.

For the GFP knockdown FACS-seq experiments, two recipient strains expressing *dcas9* were constructed: one encoding *gfp* (strain CAG78108) and the other encoding *rfp* (strain CAG78107). Each was generated by first cloning the constitutive *gfp* and *rfp* expression cassettes from pDG1731-gfp and pDG1731-rfp upstream of *frt-cat-frt* from pKD3 (*39*), integrating them into the chromosome between *yjaA* and *yjaB* using recombineering (*40*), and selecting for chloramphenicol resistance. P1 transduction (*41*) was then used to move the *gfp-frt-cat-frt* or *rfp-frt-cat-frt* cassettes into BW25113, selecting for chloramphenicol resistance. Chromosomal *dcas9* was then introduced to these strains by conjugation using a pseudo-Hfr *dcas9* donor, as described previously (*42*), where in this case the *dcas9* is expressed by the minimal synthetic promoter P*BBa_J23105* {https://parts.igem.org}, and transconjugates were selected using gentamicin and chloramphenicol.

Unless otherwise noted, all strain construction and growth assays for *E. coli* were done in LB medium and using antibiotic selection at the specified concentrations: ampicillin (100µg/ml), carbenicillin (50µg/ml), gentamicin (10µg/ml), chloramphenicol (25µg/ml).

*Bacillus subtilis* CRISPRi library construction

As in the individual CRISPRi strain construction (above), CRISPRi libraries were constructed by transforming sgRNA plasmids into the *dcas9* strain. The protocol was modified in one of two ways in order to increase the scale; we found both methods were sufficient to maintain coverage of the pooled plasmids. In one method, cells were grown in *B. subtilis* competence medium to OD600=1.5, and then incubated with plasmid DNA (300µl cells + 300ng plasmid DNA) in 96-well deep-well plates. Incubations were performed for 2hr at 37C with shaking (900RPM), after which point plates were spun down at 5000g for 10 minutes and resuspended in 2mL LB medium before plating on plates (Falcon #351058) with chloramphenicol at a density ~0.4M CFU/plate and growth overnight at 37C. A second method incubated competent cells (grown in *B. subtilis* competence medium to OD600=1.5) with plasmid DNA in culture flasks, for 2hr at 37C with shaking (900RPM), after which point cells were spun down in 50ml tubes and resuspended in 2-6ml LB before plating on chloramphenicol plates as before.

To store the transformed CRISPRi library, plates were scraped, pelleted and resuspended in S7 salts (*21*) with 15% glycerol, and stored in 500uL aliquots at -80C.

*Escherichia coli* CRISPRi library construction

Strain library construction from plasmid libraries was achieved by electroporating plasmid DNA into the recipient strains, and plating on plates (Falcon #351058) with carbenicillin and 0.2% glucose (to repress uptake of residual lactose in LB that can induce the IPTG-controlled *dcas9* in the essential gene knockdown strains) at a density ~0.4M CFU/plate and growth overnight at 37C. To store the libraries, plates were scraped, pelleted, and resuspended in 15% glycerol to be stored at -80C.

sgRNA plasmid construction

The sgRNA plasmid pJSHA77 was modified from pDG1622 to increase transformation and double-crossover efficiency. 1.5kb of DNA upstream of *amyE* was PCR amplified from *B. subtilis* 168 genomic DNA and inserted into pDG1662 by HiFi Assembly (New England Biolabs #E2621L), replacing the shorter upstream fragment of *amyE* in pDG1662. Synthetic DNA containing a transcription terminator, an sgRNA driven by P*veg* with BsaI cut sites for spacer cloning, and downstream tandem transcription terminators was purchased from IDT and cloned into the previously described pDG1662 derivative by HiFi Assembly (New England Biolabs #E2621L), generating pJSHa77.

Oligonucleotide pools containing the desired elements with flanking restriction sites and library-specific PCR adapters were obtained from Agilent Technologies (Table S8). The oligonucleotide pools were amplified by 15 cycles of PCR using Q5 polymerase (New England Biolabs #M0493S) and custom primers (Table S8). The PCR product was digested with BsaI-HFv2 (New England Biolabs #R3733) and gel purified from 10% TBE gels (Invitrogen #EC6275BOX) to remove adapter ends. pJSHA77 vector was midi-prepped (Qiagen #12143), digested with BsaI-HFv2 for 1hr, and treated with Antarctic phosphatase (New England Biolabs # M0289S), and ligation was carried out at a 1:2 (vector:insert) molar ratio using T4 DNA Ligase (New England Biolabs #M0202L). Ligations were transformed into electrocompetent cells (New England Biolabs #C3020K), recovered for 1hr at 37C in LB, and then inoculated into 100ml with carbenicillin and grown overnight. Plasmid libraries were collected by midiprep (Qiagen #12143) and analyzed by deep sequencing (Illumina MiSeq #MS-103-1002) to assess cloning efficiency and library diversity.

For individual sgRNA strains, inserts were prepared by annealing two single-stranded DNA oligos together to create the 4-base overhangs, and then annealed inserts were ligated using T4 DNA Ligase (New England Biolabs #M0202L) individually into pJSHA77 digested with BsaI-HFv2 and treated with Antarctic phosphatase (New England Biolabs # M0289S).

For the single-strain competition validation strains, pJSHA77 was first modified to incorporate a constitutively expressed P*veg-gfp* or P*veg-rfp* using HiFi Assembly (New England Biolabs #E2621L). Strains were then constructed as described above, ligating annealed-pair inserts into the modified vector after digesting with BsaI-HFv2.

sgRNA plasmid library design

Code for designing (fully matched) sgRNA spacers targeting a list of genomic loci can be found at https://github.com/traeki/sgrna_design.

Non-targeting sgRNA controls were designed by creating random 20nt sequences with a distribution of GC content similar to *B. subtilis* (~45%), and then using bowtie (*43*) to identify (and subsequently filter out) sgRNAs which aligned (allowing 3 or fewer mismatches) to other intragenic targets in the combined genomes of *E. coli* and *B. subtilis*, or any targets in *gfp* or *rfp*.

For the libraries targeting all essential genes in *B. subtilis*, multiple iterations of sgRNA library design (i.e. spacer design), construction, and analysis were used. For *B. subtilis* libraries, all presented data is from V2 library measurements, with the exception of the trimethoprim experiments which used measurements of the V1 libraries (all data in Table S3).

For the V1 libraries targeting *B. subtilis* genes we chose target genes to be all those previously identified as essential, putative essential, or low-fitness (*8*, *21*) (Table S3). For every gene in the V1 set, two non-overlapping fully complementary spacers were chosen, each targeting the non-template strand as close to the start of the ORF as possible. For each fully complementary spacer, a set of 25 spacer variants were designed and ordered: 2x the fully complementary spacer, 5x randomly chosen single-mismatches within 7 bases of the PAM, 5x randomly chosen single-mismatches 8-12 bases from the PAM, 3x randomly chosen single-mismatches 13-19 bases from the PAM (to exclude the outermost base), 10x randomly chosen double mismatches 1-19 bases from the PAM. In addition, for every gene the first three non-overlapping template-strand spacers were included.

The V2 *B. subtilis* libraries included all essential *B. subtilis* genes as well as a subset of non-essential but fitness-impacting genes (Table S3) from V1 of the library. The V2 *E. coli* libraries included a majority of genes with evidence for essentiality (Table S3) (*21*). For every gene in this set, ten non-overlapping fully complementary spacers were chosen on the non-template strand, as close to the start of the ORF as possible. For each fully complementary spacer, a set of 10 spacer variants was designed and ordered (for a total of 100 sgRNAs per gene): 1x the original fully complementary spacer, 9x single-mismatches (Fig. S1). Single-mismatches were chosen using the following criteria: all possible single-mismatch variants were evaluated by the trained linear model for a predicted sgRNA activity {https://github.com/traeki/mismatch_crispri, train_linear_model.py and choose_guides.py}. These predicted sgRNA activities were categorized into five bins: <10%, >90%, and three equally sized bins between 10% and 90% predicted sgRNA activity. Three sgRNAs were chosen from each of the middle three bins. For the design of all libraries using this strategy, a preliminary version of the linear model was used.

The compact libraries with 11 sgRNAs per gene were selected as above, with the following modifications for both species: for each gene 2x fully complementary sgRNAs were chosen, and 9x single-mismatch variants were selected from among all possible single-mismatch variants of each, using a binning strategy as described above (Fig. S1). For *E. coli*, also as described above, the bins were generated using predicted sgRNA activity. For *B. subtilis* the bins were instead generated using the measured relative fitness values from the V1 experiment, and the selected sgRNAs were therefore a subset of those used in the V1 library.

The *dfrA, gfp,* and *rfp* V1 comprehensive libraries (used in the trimethoprim experiment and all FACS-seq experiments) were designed analogous to the V1 essential gene libraries, with 100 sgRNAs per target: 4x the original fully complementary spacer, 20x randomly chosen single-mismatches within 7 bases of the PAM, 15x randomly chosen single-mismatches 8-12 bases from the PAM, 12x randomly chosen single-mismatches 13-20 bases from the PAM, and 49x randomly chosen double mismatches 1-20 bases from the PAM (Fig. S1).

For the V2 comprehensive libraries targeting *dfrA*, *murAA*, *folA*, or *murA*, we designed all possible non-template spacers, each with all possible single-mismatches, for a total of 60x mismatch variants per fully complementary sgRNA.

### Methods

#### Relative fitness experimental details

Glycerol stocks of the *B. subtilis* essential-gene library (V1 or V2), the *dfrA* and *murAA* libraries (V1 or V2), and the library of non-targeting control sgRNAs were fully thawed, mixed, and inoculated into 150 mL cultures of LB at a combined OD600 of 0.01 (5% control, 75% essential-gene library, 10% *dfrA* library, 10% *murAA* library). This culture was allowed to grow to OD600 0.1, at which point the culture was back-diluted to OD600 0.01 in fresh 150 mL culture of LB + 1% xylose. This culture was then grown to OD600 0.3 (~5 doublings), back-diluted to OD600 0.01 in LB + 1% xylose, and grown to OD600 0.3 (total ~10 doublings). Samples were collected a) immediately before back dilution into xylose and b) after the final growth phase, ~10 doublings apart (Fig. 2A). The trimethoprim experiments were carried out in an identical manner, except that both 1% xylose and trimethoprim (0 ng/mL, 15ng/mL, or 30ng/mL) were added from the first back-dilution and maintained throughout growth. Concentrations of trimethoprim were chosen such that wildtype growth rate was unaffected.

Fitness experiments for the *E. coli* V2 libraries were carried out in an identical manner to the *B. subtilis* fitness experiments with the following exceptions: all growth occurred in the presence of ampicillin, and induction was achieved with 1mM IPTG instead of 1% xylose.

For both *B. subtilis* and *E. coli*, compact library experiments were carried out in an identical manner as the larger scale fitness experiments above, save that the volume of cultures was 15mL, and only compact libraries and non-targeting control libraries were mixed together (90% compact library, 10% controls).

At the desired time points, *B. subtilis* cultures were collected (1ml) by pelleting (9000xg 2min) and genomic DNA was extracted using the DNeasy Blood & Tissue kit (Qiagen #69506) with the recommended Gram-positive pre-treatment and RNAse A treatment. For the *E. coli* fitness experiments, *E. coli* cultures were collected (4ml) by pelleting (20000xg 2min) and plasmid DNA was extracted using the QIAprep Spin miniprep kit (Qiagen #27106). sgRNA spacer sequences were amplified from gDNA or plasmid DNA using Q5 polymerase (New England Biolabs #M0493S) for 14x cycles using custom primers containing TruSeq adapters and indices (Table S8), followed by gel-purification from 8% TBE gels (Invitrogen #EC62152BOX), and sequencing on HiSeq 4000 with single-end 50bp reads using a custom sequencing primer (UCSF Center for Advanced Technology).

#### Relative fitness analysis

Raw FASTQ files were aligned to the library oligos and counted using {https://github.com/traeki/mismatch_crispri, count_guides.py}, and relative fitness was calculated using {https://github.com/traeki/mismatch_crispri, compute_gammas.py and gamma_to_relfit.py}. For each strain ($x$) with at least 100 counts at $t_0$ we calculate the relative fitness $F(x)$ according to:

$$F(x) = \frac{log2\,\frac{r_{wt}(t_0) * r_x(t_{10})}{r_{wt}(t_{10}) * r_x(t_0)}}{g_{wt}} + 1$$

where $r_x(t_i)$ is the fraction of strain $X$ in the population at time $i$ and $g_{wt}$ is the number of generations of wildtype growth in the experiment. A derivation of this equation can be found in (*3*) and (*16*). In our experiments, $g_{wt}$ is calculated from the OD measurements of the culture, and $r_{wt}(t_i)$ is calculated as the median of 1000 non-targeting control sgRNAs from that sample. For strains with at least 100 counts at $t_0$ and 0 counts at $t_{10},$ we set:

$$log2\,\frac{r_x(t_{10})}{r_x(t_0)} = 0$$

Finally, the relative fitness measurements of each sgRNA were averaged across samples (*B. subtilis* experiments: 6 replicates, *E. coli* experiments: 4 replicates) to calculate the final relative fitness value and standard deviation (Table S3).

Relative fitness validation

To validate the practice of using pooled growth measurements as an approximation of relative fitness, we also measured the relative fitness of individual *dfrA* knockdown strains grown in the presence of a wildtype strain. For each *dfrA* sgRNA, the spacer was cloned separately into pJSHA77-gfp and pJSHA77-rfp, each transformed into the *dcas9* strain, and then competed against a wildtype constitutively expressing the opposite fluorophore (i.e. strains with a *dfrA* sgRNA and expressing *gfp* were competed against a wildtype expressing *rfp*). Strains were mixed at a starting OD600 of 0.01 in 300µL of LB in four replicate wells of a 96-well deep-well plate, covered with a breathable film, and grown shaking at 900 RPM at 37C. Cells were diluted to OD600 0.01 in fresh LB with 1% xylose and grown again (900 RPM, 37C) to OD600 0.3. Immediately after each back-dilution (and at end of experiment) the previous plate was fixed with 50µL of 37% formaldehyde per well, incubated for 10min at room temperature, and quenched with 50µL of 2.5 M glycine. The quenched reaction was diluted 1:20 into 1X PBS before measurement by flow cytometry (LSRII, BD Biosciences) using the blue laser (488 nm) and the FITC detector (530/30 nm) for GFP detection, and the yellow/green laser (561 nm) and the PE-Texas Red detector (610/20 nm) for RFP detection. Data for at least 20,000 cells were collected, and thresholds based on control wells were used to define the GFP+ and RFP+ populations to determine the ratio of each population in each sample using FlowJo (FlowJo, LLC). All calculated relative fitness measurements from this validation experiment are provided in Table S3.

FACS-seq experimental details

Three separate strain libraries were constructed and mixed together for use in the sorting experiments: a *gfp*$^+$ strain with the *gfp*-targeting sgRNA library (mismatch-GFP), a *gfp*$^+$ strain with the non-targeting sgRNA control library ("high-GFP" or "control sgRNA" in figure), and a *gfp*$^-$ strain with the *rfp*-targeting sgRNA library ("no-GFP" or "dark control") (Fig. 1A). Glycerol stocks of each library were fully thawed, inoculated into replicate 12.5ml cultures of LB (*B. subtilis*) or LB with ampicillin (*E. coli*) at 0.01 OD600, and allowed to grow for 2.5-3hr. Then cultures were back-diluted to 0.01 OD600 in LB with 1% xylose (*B. subtilis*) or LB with ampicillin (*E. coli*) and grown for 2.5hr. Immediately before sorting the cultures were mixed at a ratio reflecting the overall diversities of their libraries (40% mismatch-GFP, 40% low-GFP, 20% high-GFP), and then the mixture was diluted 1:10 in PBS at room temperature (*B. subtilis*) or on ice (*E. coli*).

Sorting was done on the mixed cultures using a BD FACSAria II (Laboratory for Cell Analysis in Helen Diller Family Comprehensive Cancer Center at UCSF), using the blue laser (488 nm) and the FITC detector (530/30 nm), and at a flow rate of 5 and collecting for 20min total. Post-sorting the collected bins were filtered using either cellulose nitrate membranes with 0.2um pore (Thermo Scientific #145-0020) or mixed cellulose esters 0.22um pore disc filters (MF-Millipore #GSWP02500) on a glass filtration apparatus. Filters were resuspended in 9ml LB (*B. subtilis*) or LB with ampicillin (*E. coli*) by vortexing at max speed for 30s, then split into two outgrowth cultures and grown overnight in 4ml LB (*B. subtilis*) or LB with ampicillin (*E. coli*). A portion of the input mixed sample (i.e. pre-sorting) was treated similarly and grown overnight. DNA was extracted from each outgrowth culture separately and analyzed by deep sequencing as described above.

<u>FACS-seq analysis</u>

For each species, two biological replicates (i.e. cultures starting from unique glycerol stocks) were sorted by FACS, and from each biological replicate's 4 bins (plus unsorted mixture) two technical replicates (i.e. two overnight outgrowth cultures from which DNA was extracted) were sequenced. Library spacers were counted in each sequenced sample, normalized to the sample's total number of spacers counted, and technical replicate normalized counts were added together. For each biological replicate, the sorted bins were further normalized with respect to the mixed (i.e. pre-sorting) sample in the following manner: a linear model was used to determine the appropriate weights for each bin in order to recapitulate the mixed sample, and those weights were applied as scaling factors for all read counts from the given bin. This normalization was essential to correct for sequencing depth and cell number differences between bins. Briefly, we used the sklearn package (sklearn.linear_model) in Python and applied it to the mixed sample after removing from it the top and bottom 5$^{th}$ percentiles.

We sought to define a metric for enrichment in the GFP-high bins vs. the GFP-low bins that would be similar in scale to relative fitness. We define an enrichment ratio (ER) for each sgRNA as:

$$ER = \frac{3}{3} n.norm_{Bin4} + \frac{2}{3} * \ n.norm_{Bin3} \ \frac{1}{3} * \ n.norm_{Bin2} + \frac{0}{3} * \ n.norm_{Bin1}$$

where $n.norm_{Bin\ i}$ is the normalized counts in Bin *i*, and Bin1 has the lowest GFP fluorescence while Bin4 has the highest. By this metric, values close to 1 have the highest GFP fluorescence (or weakest sgRNA activity) and values <1 have lower GFP fluorescence (or stronger sgRNA activity). Enrichment scores were normalized on a per experiment basis by subtracting the mean enrichment score of the "dark controls" and dividing by the mean enrichment score of the "high-GFP" strains. The resulting scores for each sgRNA (called the "FACS-seq score" in the main text) are available in Table S1.

<u>FACS-seq validation</u>

To validate our sorting procedure and the relationship between the calculated FACS-seq score and the fluorescence of a single strain, we randomly isolated 9 strains from the *E. coli* GFP knockdown library and analyzed them by flow cytometry to quantify knockdown relative to a non-targeting sgRNA (Fig. S2E-H). Strains were grown in deep 96-well plates in 300ul LB overnight, diluted back and grown to ~0.4 OD600 before measurement. Briefly, data was collected on a LSRII flow cytometer (BD Biosciences) using the blue laser (488 nm) and the FITC detector (530/30 nm). Data for at least 20,000 cells were collected, and median fluorescence values were extracted using FlowJo (FlowJo, LLC). Data from representative samples were plotted as histograms using FlowJo to confirm that single-cell fluorescence was unimodal within the population (Fig. S2E-H). sgRNA plasmids were miniprepped (Qiagen #27106) from each library isolate and Sanger sequenced to ascertain their identity in the library experiment. To assay the behavior of the same sgRNAs in *B. subtilis,* the miniprepped plasmid was transformed into *B. subtilis* as described above, double-crossover events were verified by streaking on starch plates, and the strains were analyzed by flow cytometry as described above. All relative fluorescence measurements are provided in Table S1 and plotted in Fig. S2A.

Predicted sgRNA activity validation

To validate the linear model's ability to predict sgRNA activity based on sgRNA sequence, we measured the knockdown of a *murAA-gfp* transcriptional fusion in a *B. subtilis* strain that was complemented by a non-targeted copy of *murAA*. These strains also expressed a chromosomal *rfp* that allowed for calculation of the GFP/RFP ratio on a per cell basis. Strains were grown as described above (FACS-seq validation), with the exception that *dcas9* was induced using 1% xylose after dilution. Data was collected on a LSRII flow cytometer (BD Biosciences) using the blue laser (488 nm) and the FITC detector (530/30 nm) for GFP detection, and the yellow/green laser (561 nm) and the PE-Texas Red detector (610/20 nm) for RFP detection. Data for at least 20,000 cells were collected, and the per-cell GFP/RFP ratios as well as the population median GFP/RFP ratios were extracted using FlowJo (FlowJo, LLC). Relative knockdown was normalized to a *murAA-gfp* strain lacking a sgRNA, after first subtracting the background GFP fluorescence from a non-fluorescent *B. subtilis* strain. Relative GFP fluorescence measurements are provided in Table S10.

Linear model of singly mismatched sgRNA efficacy

Having measured the ability of ~1,600 singly mismatched sgRNAs to knockdown GFP expression, we sought to build a model to predict the effect of mismatches on sgRNA efficacy. Since an enrichment score of 1 represent maximal GFP fluorescence, and a score of 0 represents no GFP fluorescence, we define knockdown for each sgRNA as:

$$knockdown_{sgRNA} = 1 - FACS.seq\ score$$

We then normalized the ability of each mismatched sgRNA to knockdown GFP compared to its equivalent fully complementary sgRNA using the equation below:

$$sgRNA\ activity_{singly\ mismatched\ sgRNA} = \frac{knockdown_{singly\ mismatched\ sgRNA}}{knockdown_{fully\ complementary\ sgRNA}}$$

We next built a model that fit the activity of each sgRNA using the position of the mismatch (from 0 to 19, with 19 being PAM proximal, one hot encoded), the transition of the mismatch (from X to Y, one hot encoded), and the GC% of the fully complementary sgRNA.

Mismatched sgRNAs were excluded from the analysis if they were variants of fully complementary sgRNAs with less than 0.5 knockdown (as described above). The parameters from this model trained on *E. coli, B. subtilis,* or species-averaged per sgRNA activity are presented in Table S2 and Fig. S10, the raw data in Table S1.

Expression-fitness relationship analysis

In order to quantitatively assess the expression-fitness relationship of genes targeted by the V2 *E. coli* and *B. subtilis* libraries, we developed a per gene pipeline, described below.

1. In general, fully complementary sgRNAs targeting the same gene had similar fitness effects (Fig. S11 and supplementary text 1), suggesting that all fully complementary sgRNAs induce a similar level of knockdown. We identified outlier sgRNAs that were significantly less effective at inducing a fitness defect (and therefore were likely to be ineffective at knocking down their target) by comparing the distribution of fitness values for each series (series = the fully complementary sgRNA and its 9 singly mismatched variants) to the fitness distribution of the remaining series targeting the same gene. Using a two-sided t-test, we assessed whether the distribution of their relative fitness values was significantly different ($p < 0.05$) from the relative fitness distribution of the remaining sgRNAs targeting the gene. If their distribution was significantly different and their mean relative fitness was higher than the other sgRNAs targeting the same gene, we surmised that the fully matched sgRNA was likely not functional and excluded its series from further analysis.

2. We next predicted the sgRNA activity of all sgRNAs using the model of sgRNA efficacy described above trained on the two species averaged GFP data also described above. Consistent with the definition of sgRNA activity above, fully complementary sgRNAs were assigned a sgRNA activity of 1.

3. We binned sgRNAs that passed our filter (Step 1) based on their predicted sgRNA activity (bin width = 0.2, bin spacing = 0.05, for a total of 17 bins), and within each bin we calculated the median relative fitness. A fully healthy (relative fitness = 1, predicted sgRNA activity = 0) pseudocount was included for each gene. Per sgRNA and per gene patterns are shown in Fig. S8 (*E. coli*) and Fig. S9 (*B. subtilis*). Per gene bin medians for essential genes can be found in Tables S5.

Per gene bin medians were used in all analyses of gene expression-fitness relationship similarity.

Gene expression-fitness relationship clustering and enrichment analysis

To determine whether per gene expression-fitness curves were biologically meaningful, we clustered the bin medians (described above) for all essential gene in *E. coli* and *B. subtilis* into 9 clusters using k-means with 10,000 random restarts. Functional enrichment within clusters was calculated for COG categories, GO biological process terms, and KEGG terms using the hypergeometric test. Only p-values with Bonferroni corrected ($p < 0.05$) are shown in Table S6.

Gene similarity comparisons

To determine whether the expression-fitness relationships of genes within COG categories, GO biological process, or KEGG categories were more similar to each other than to those of other genes we first calculated pairwise Euclidean distances between the expression-fitness relationships of all essential genes within each species. We then used a two-sided t-test to compare the distances between genes within each category to the distances between those genes and genes in different categories. We accounted for CRISPRi polarity due to operon structure by excluding any distances between genes within the same operon (defined as two genes in the same direction <50bp apart) from both the "inside category" and the "outside category" set.

To determine whether the expression-fitness relationships of homologous genes were more similar to each other than to those of other genes in the opposing organism, we calculated the pairwise Euclidean distance between the expression-fitness relationships of all essential genes that have essential homologs in both *E. coli* and *B. subtilis* (n = 155, as defined in Koo et at., 2017). We next used a two-sided t-test to determine if the distance between homologs was, on average, different from the overall distribution of distances between these 155 genes (i.e. when one gene from one species is compared to the 154 genes in the opposing species). To determine which pairs of homologs were significantly dissimilar, for each gene pair (including homologs), we calculated how many cross-species comparisons involving either gene were more similar than the comparison in question. We compared this number in homologs and non-homologs to calculate a FDR.

**Supplementary Text**

1. Quantifying similarity between fully complementary guides targeting the same gene

Our *gfp* based model predicts the activity of singly mismatched sgRNAs relative to the activity of the fully complementary sgRNA from which they are derived. To use this relative sgRNA activity as a proxy for absolute activity, fully complementary sgRNAs targeting the same gene should have the same activity. Since we cannot easily measure the sgRNA activity directly when targeting endogenous essential genes, we reasoned that we could validate this assumption by comparing fitness effect of fully complementary sgRNAs targeting the same gene. To do so, we compared the total sum of squares (totalSS) of fully complementary sgRNAs to within gene the sum of squares (withinSS). In *E. coli*, the withinSS accounted for 26.7% of the totalSS and in *B. subtilis* the withinSS accounted for 18.6% of the totals. This suggests that fully complementary sgRNAs targeting the same gene are substantially more similar with regards to their fitness outcomes than fully complementary sgRNAs as a whole, and supports the assumption that fully complementary sgRNAs targeting the same gene have similar levels of activity.

2. Detection limits of relative fitness measurements

Our relative fitness experiments seek to quantify the number of doublings each strain experiences during the course of the experiment, relative to the number of doublings a wild-type (or a non-targeting sgRNA control) strain experiences during this time. To do so, we measure the bulk growth of the population, and quantify the relative abundance of each strain at the start and end of each experiment *via* next-generation sequencing. Changes in the relative abundance of a strain are determined by the growth rate of the individual strain relative to the population as a hole. For example, there is a $2^{10} \sim 1{,}000$-fold increase in the number of cells during a 10 doubling experiment. Therefore, cells that do not divide (but remain intact) will experience a 1,000-fold decrease in relative abundance.

Our ability to measure the relative abundance of strains is constrained by sequencing depth. Assuming an equal number of reads at the start and end of the experiment, measurement of a 1,000 fold decrease in relative abundance requires that a strain have at least 1,000 reads at the start of the experiment. A poorly represented strain (e.g. 50 read counts at the start of the experiment) cannot decrease 1,000-fold and be meaningfully measured.

Previously reported pooled fitness experiments of CRISPRi libraries in *E. coli* prioritized sensitivity to slight growth defects over quantifying the extent of a strong fitness defect (Fig. S6D-E). To do so, these experiments were run for many generations (15+) and were sequenced with relatively less depth (median counts ~100). This limited their ability to quantify strong fitness effects. In contrast, this study prioritized quantification of the full range of possible fitness outcomes. As a result, our experiments were run for 10 generations and deeply sequenced (median counts > 1,000), allowing us to quantify a broad range of fitness defects.

Many strains were abundant enough at the start of the experiment to allow accurate quantification of decreases greater than $2^{10} \sim 1{,}000$-fold. These events (relative fitness < 0) represent active depletion from the pool.

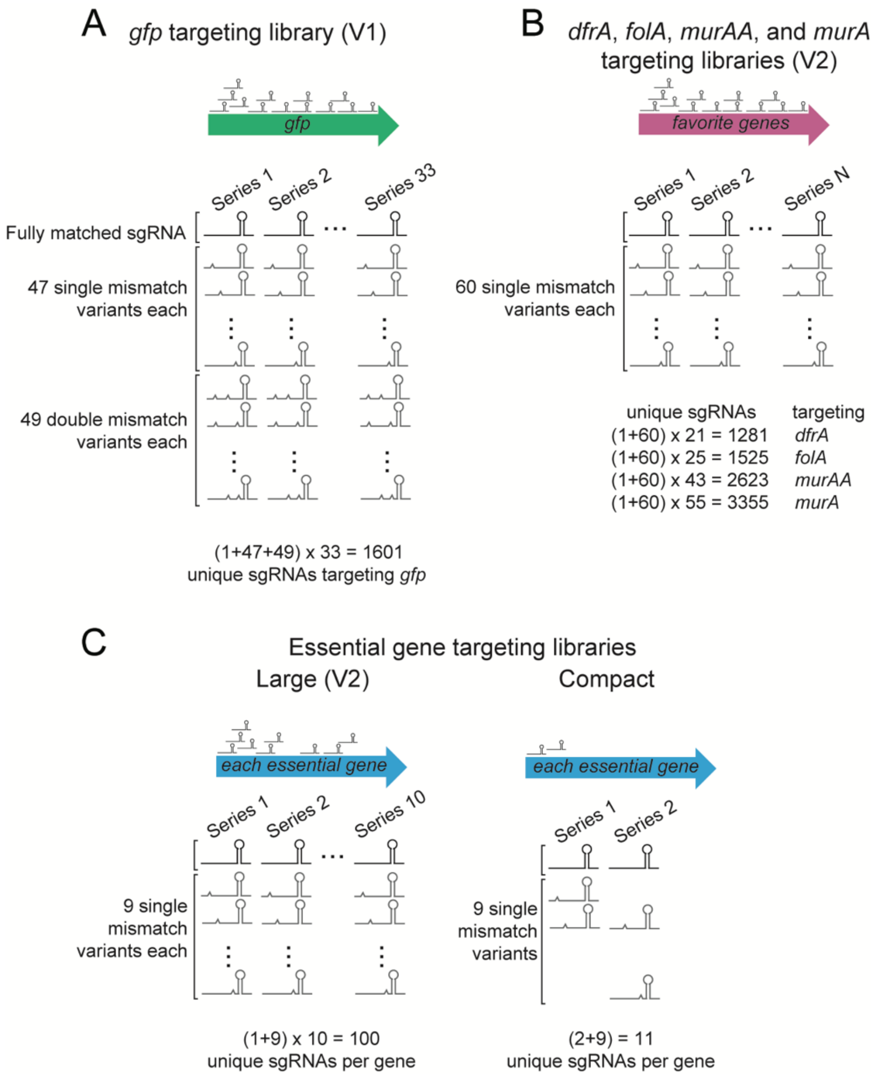


**Fig. S1.** Design of mismatched sgRNA libraries targeting (A) *gfp*, (B) comprehensive libraries targeting *dfrA*, *folA*, *murAA*, and *murA*, and (C) each essential gene for the large libraries and the compact libraries. The breakdown of single mismatch variants per series and the total unique sgRNAs per gene are shown for each.

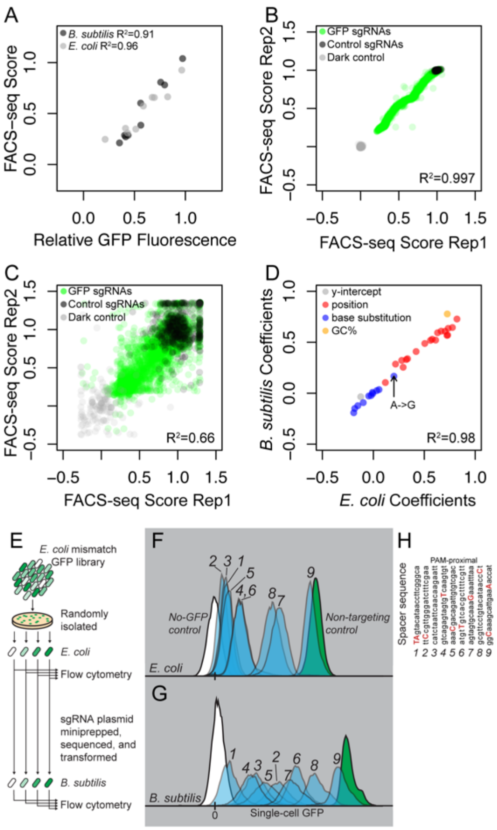
A
B. subtilis R²=0.91
E. coli R²=0.96
FACS–seq Score
Relative GFP Fluorescence
B
GFP sgRNAs
Control sgRNAs
Dark control
R²=0.997
FACS-seq Score Rep2
FACS-seq Score Rep1
C
GFP sgRNAs
Control sgRNAs
Dark control
R²=0.66
FACS-seq Score Rep2
FACS-seq Score Rep1
D
y-intercept
position
base substitution
GC%
A->G
R²=0.98
B. subtilis Coefficients
E. coli Coefficients
E
E. coli mismatch GFP library
Randomly isolated
E. coli
Flow cytometry
sgRNA plasmid miniprepped, sequenced, and transformed
B. subtilis
Flow cytometry
F
No-GFP control
Non-targeting control
E. coli
G
B. subtilis
Single-cell GFP
H
PAM-proximal
Spacer sequence

**Fig. S2.** Details related to the linear model, FACS-seq data, and its validation. (A) Mismatched sgRNA efficacy measured individually (relative GFP fluorescence) in either *E. coli* or *B. subtilis* compared to their FACS-seq score from measurements done in the same species. Relative fluorescence is the median GFP single-cell fluorescence, normalized as a fraction of non-targeted control. (B) FACS-seq scores for all sgRNAs comparing two biological replicates in *B. subtilis*. (C) FACS-seq scores for all sgRNAs comparing two biological replicates in *E. coli*. Increased noise in *E. coli* likely reflects variation in sgRNA plasmid copy number at the time of DNA extraction and/or sequence-based *E. coli* specific effects on sgRNA efficacy (*20*). (D) Coefficients of each variable from linear models trained on FACS-seq data from either *E. coli* or *B. subtilis*. The A to G transition discussed in the main text is highlighted. y_int is the y-intercept; GC is the GC% of the fully complementary sgRNA spacer. (E) Schematic describing the isolation of random singly mismatched sgRNA strains from the *E. coli gfp* library, their analysis by flow cytometry, the introduction of the same sgRNA plasmids into *B. subtilis*, and their analysis by flow cytometry. (F-G) The distribution of single-cell GFP fluorescence values for strains of *E. coli* (G) or *B. subtilis* (F). (H) The sequences of the spacers indicated in (F) and (G), with mismatched bases highlighted in red.

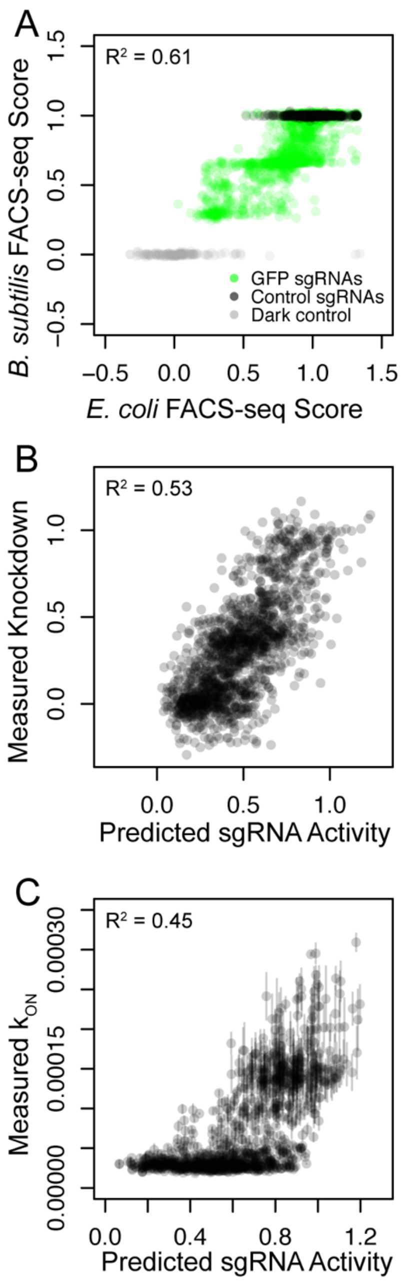


**Fig. S3.** Doubly mismatched sgRNAs are accurately predicted as the combined independent effects of singly mismatched sgRNAs. (A) FACS-seq enrichment scores (average of 2 biological replicates) for each doubly mismatched sgRNA targeting *gfp* in *B. subtilis* and *E. coli*. (B) The predictions of the linear model for doubly mismatched sgRNA efficacy, treating each mismatch as independently affecting sgRNA efficacy, compared to the doubly mismatched sgRNAs' measured *gfp* knockdown efficacies (two species averages). (C) The predictions of the linear model for doubly mismatched sgRNAs compared to the measured doubly mismatched sgRNA association rates ($k_{ON}$) in vitro (*14*).

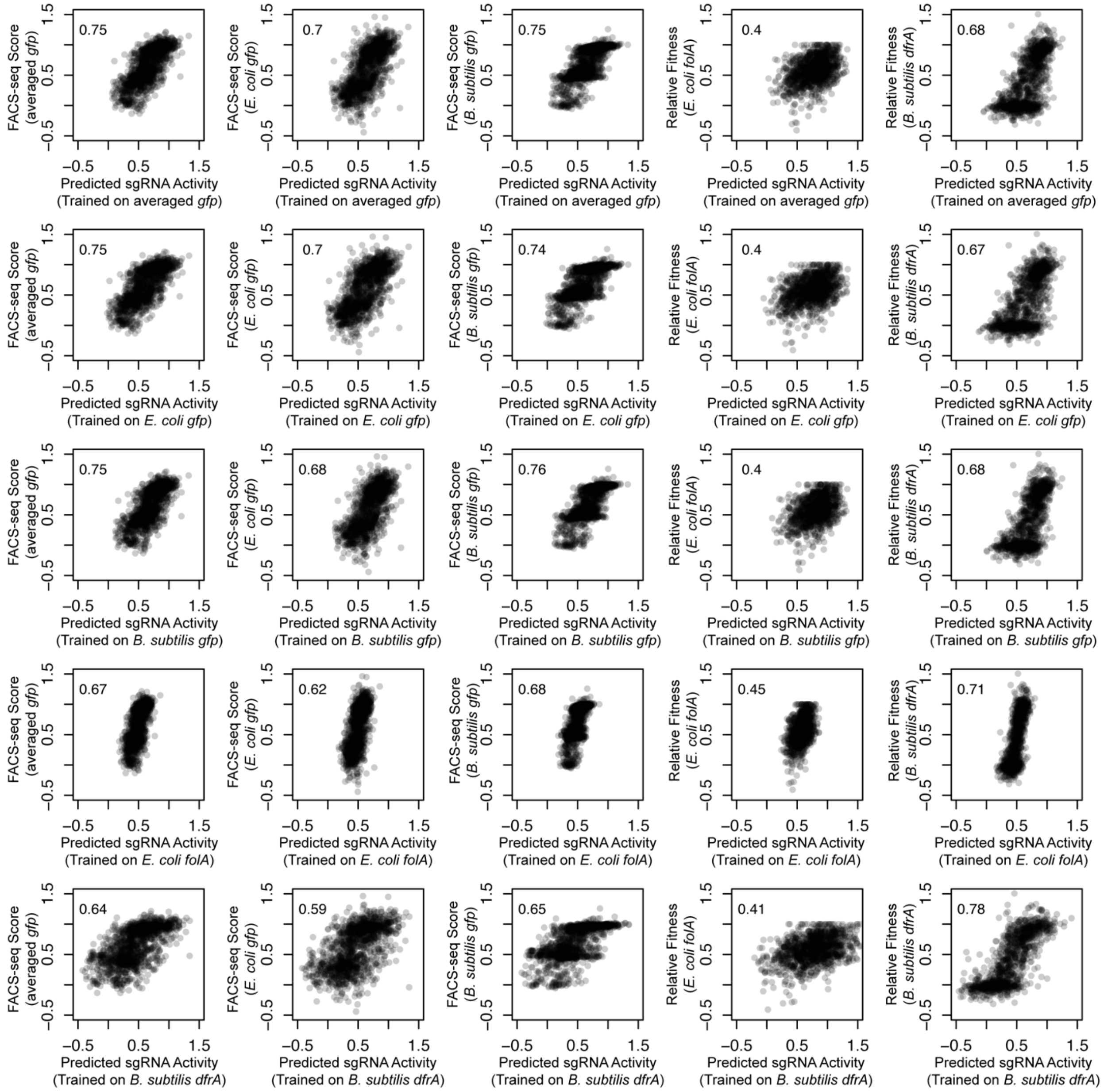


**Fig. S4.** Linear models of singly mismatched sgRNA efficacy trained on FACS-seq or relative fitness data from *E. coli, B. subtilis,* or the average of both (averaged *gfp*) retain a majority of their predictive power on other singly mismatched sgRNA datasets. Each panel compares the predictions of a linear model trained on the specified dataset to the measured efficacy (relative fitness or FACS-seq score) of sgRNAs in the other specified dataset, with the pearson correlation coefficient shown in the inset. The datasets used and evaluated are, in order from top-bottom and left-to-right: species averaged *gfp, E. coli gfp*, *B. subtilis gfp*, *E. coli folA*, and *B. subtilis dfrA*.

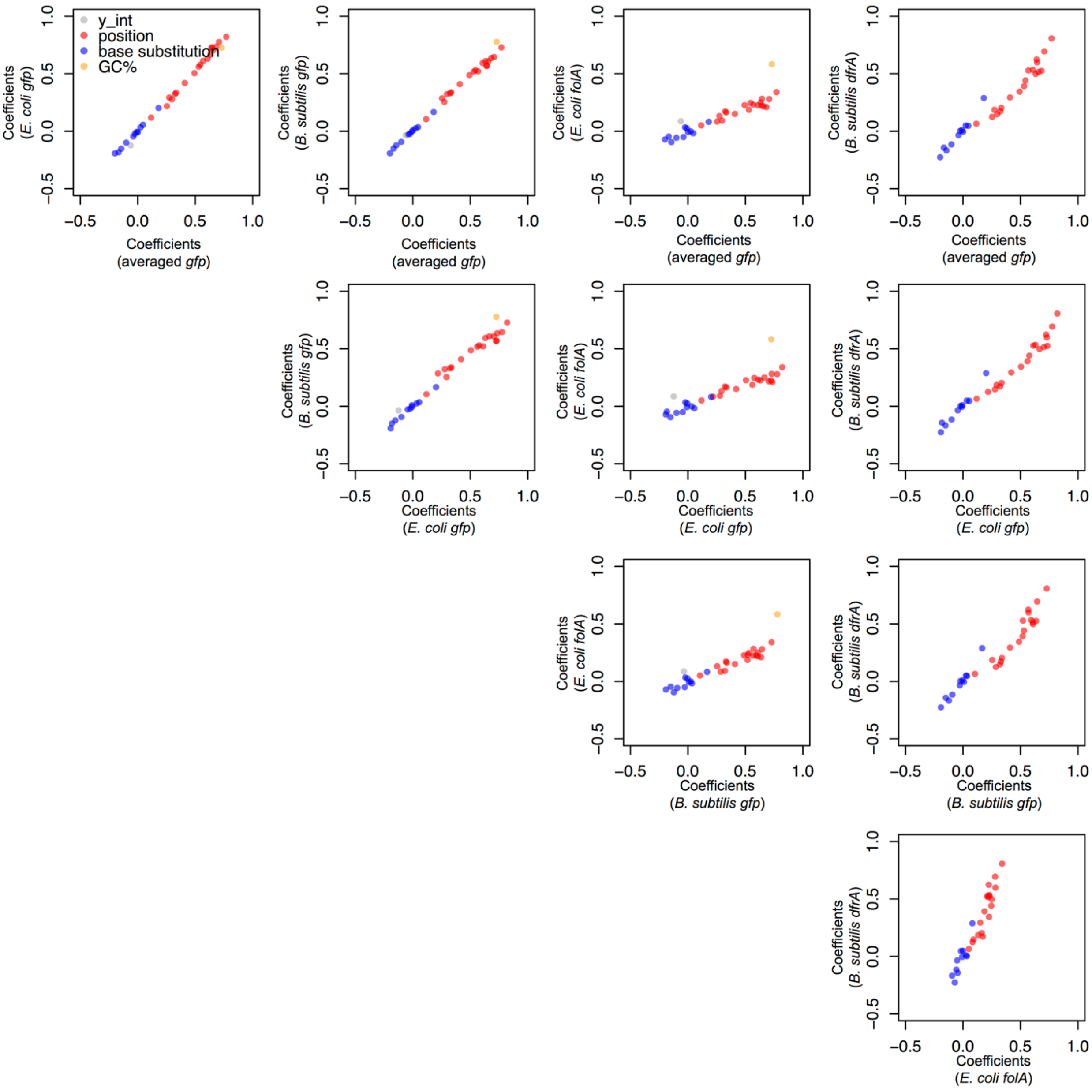


**Fig. S5.** Parameters of linear models of singly mismatched sgRNA efficacy trained on FACS-seq or relative fitness data from either *E. coli* or *B. subtilis* have strongly correlated coefficient values. Each panel compares the coefficient values from the linear models trained on the two specified datasets.

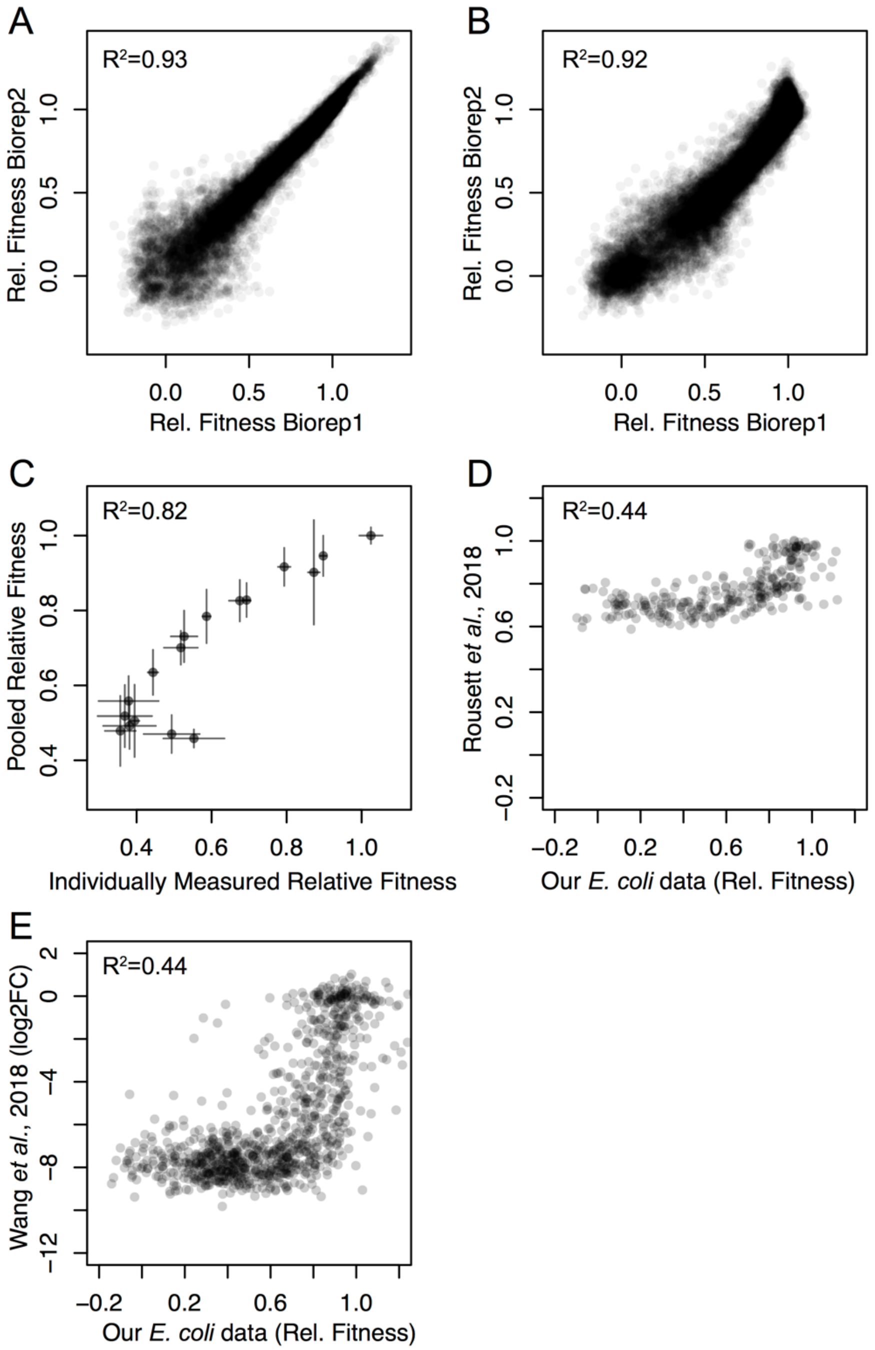


**Fig. S6.** Relative fitness measurements are reproducible, orthogonally validated, and capture a large dynamic range. (A-B) Relative fitness measurements from two biological replicates in *E. coli* (A), and *B. subtilis* (B). (C) Relative fitness from pooled experiment compared to a relative fitness metric from competing individual *dfrA*-targetomg CRISPRi strains against a fluorescently labeled wildtype and enumerating their relative abundance by flow cytometry before and after 10 doublings. (D & E) Per sgRNA relative fitness compared to previously reported fitness measurements (*19*) and (*18*) showing the increased dynamic range of our measurements. The minimum quantifiable relative fitness can be approximated by the log2(median per sgRNA read count) divided by number of generations of growth. In (*18*), median read count per sgRNA was ~100, and strains were grown for ~15 generations; therefore, relative fitness below ~0.6 is not resolvable. Similarly, in (*19*), median read count per sgRNA was >200 (~17 million total counts, 92,919 elements), and strains were grown for ~17 generations; therefore, relative fitness below 0.6 is not resolvable. See Supplementary Text 2 for further discussion.

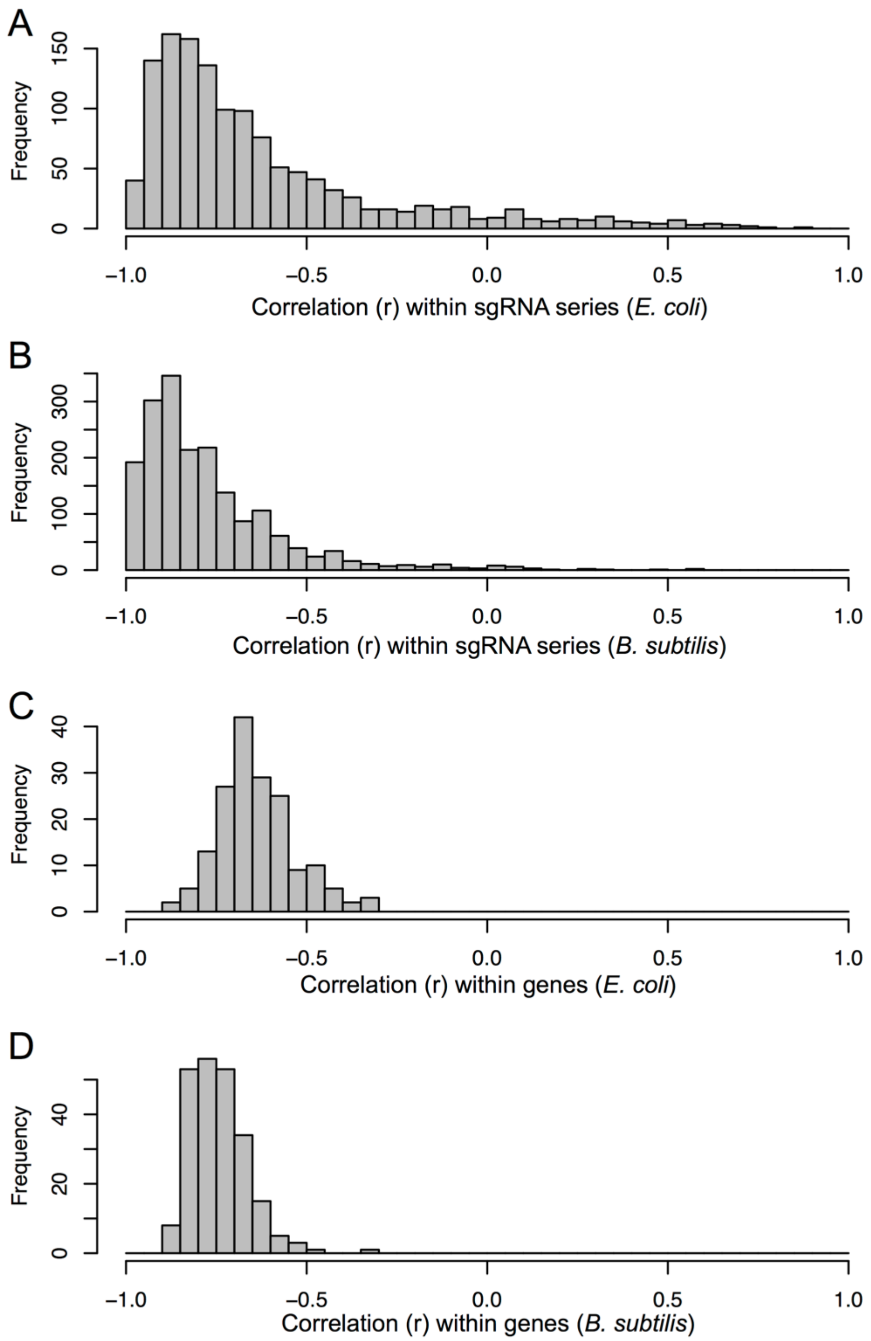


**Fig. S7.** Singly mismatched sgRNAs targeting essential genes are strongly and negatively correlated within sgRNA series and within genes. (A-B) Distribution of per sgRNA series correlations (r) for sgRNAs targeting genes in *E. coli* (A) and *B. subtilis* (B). (C-D) Distribution of per gene correlations (r) for sgRNAs targeting genes in *E. coli* (C) and *B. subtilis* (D).

**Fig. S8.** All *E. coli* per gene knockdown-fitness curves, attached as separate *.pdf.

**Fig. S9.** All *B. subtilis* per gene knockdown-fitness curves, attached as separate *.pdf.

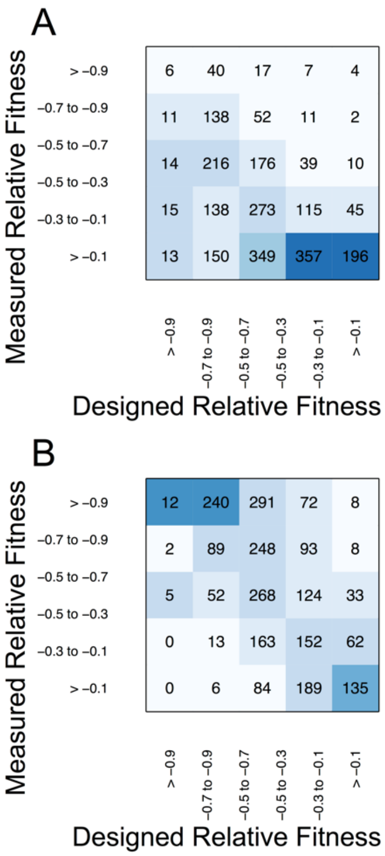


**Fig. S10.** Design constraints and measured efficacy of singly mismatched sgRNAs in the compact libraries. (A) Confusion matrix showing the singly mismatched sgRNAs selected for the compact library based on their predicted knockdown vs. their measured relative fitness in *E. coli*. Designed relative fitness shows the predicted knockdown normalized by the strongest efficacy sgRNA's relative fitness. (B) Same as (A) for the compact library in *B. subtilis*.

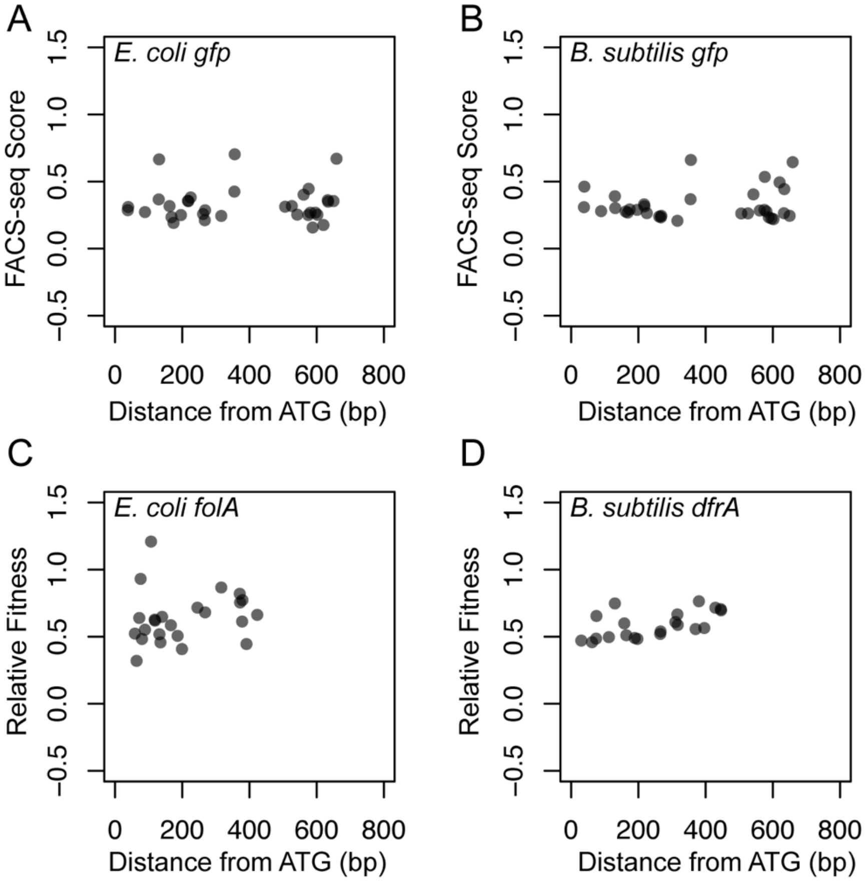


**Fig. S11.** Fully complementary sgRNA efficacy is not significantly correlated with distance within the open reading frame in *E. coli* or in *B. subtilis*. (A-B) sgRNA distance from *gfp* ATG compared to its FACS-seq score in *E. coli* (A) and *B. subtilis* (B). (C) sgRNA distance from *folA* ATG compared to its impact on fitness in *E. coli*. (D) sgRNA distance from *dfrA* ATG compared to its impact on fitness in *B. subtilis*.

**Table S1.**

FACS-seq data (enrichment ratios, training features for model, and relative *gfp* knockdown).

**Table S2.**

Linear model parameters and weights from trained models.

**Table S3.**

Fitness Data for Essential Gene sgRNA guides from Competitive Pooled-Growth Experiments.

**Table S4.**

Lysis Phenotypes for *B. subtilis* and *E. coli*

**Table S5.**

Ontological annotations and phenotype clustering data for essential genes.

**Table S6.**

Statistically significant functional enrichments from k=9 kmeans clustering in *E. coli* and *B. subtilis.*

**Table S7.**

Similarity of the expression-fitness relationship between essential homologous genes in *E. coli* and *B. subtilis.*

**Table S8.**
List of primers used in this study.

| oligo name | oligo sequence | brief description |
|---|---|---|
| agilent.pool.example | nnnnnnnnnnnnnnnnnnnnGTTTAGAGAC CCCAGTTCATTTCTTAGGG | Agilent oligo pool (example Library A), |
| agilent.amplify.example.F | atgCAAGCCTATTTTGACCAGCTCGATCA TTTTGCCCCTGGTTCTTggtc | oligos from Agilent oligo pool (example |
| agilent.amplify.example.R | GCGTTCGTTATGAAGGCTCAAAATCCTC CCCTAAGAAATGAACTGGggtc | oligos from Agilent oligo pool (example |
| JSH.E09.a77.mono.3p.1.F | [illegible] TACACGATCGGAAGAGCACACGTC TGAACTCCAGTCACATCACGCGACT CGGTGCCACTTTTTC | [illegible] amplify the spacer sequence from pJSHA77 for deep |
| JSH.D49.a77.mono.5p.R | caagcagaagacggcatacgaGATCCAGAA AAGACACCCGAAAGG | the spacer sequence from pJSHA77 for deep |
| JSH.D59.a77.mono.3p.dseq | [illegible] CGGACTAGCCTTATTTTAACTTGCT ATTTCTAG | [illegible] sequencing libraries generated from |
| murAA_bsu_F | AGAATACCATGGAAAAAATCATC | murAA into pJMP3 |
| murAA_bsu_R | AGAAAC | murAA into pJMP3 |
| murAA_m1_F | GCGCTTCTG | murAA into pJMP3 |
| murAA_m1_R | GGGTGAG | murAA into pJMP3 |
| murAA_m2_F | GAAGG | murAA into pJMP3 |
| murAA_m2_R | CGCATG | murAA into pJMP3 |
| sacArfp_5L | ATCTGGTCAATGGCGAATGTC | flanking sequence to replace sacA with Pveg- |
| sacArfp_5R | CGTTAATAAGGCTAGCTGTCGACATGTA TCCAAAATCGTCCAGCC | flanking sequence to replace sacA with Pveg- |
| sacArfp_3L | TAGA GAGAGCACAGATACGGCG ATCCGACCATTGACTGCCAC | flanking sequence to replace sacA with Pveg- |
| sacArfp_3R | CATCGGCAGC CTGTTTATTT G | flanking sequence to replace sacA with Pveg- |
| murAA_gfp_5pL | TTCGTAAAATGCGTGCGTCTG | ORF region to construct murAA-gfp |
| murAA_gfp_5pR | GTAGTTCCTCCTTATGTACTAGTAGTTTA TGCATTTAAGTCAGAAACGAC | ORF region to construct murAA-gfp |
| murAA_gfp_3pL | GAGAGCACAGATACGGCGATCAGTATG TCACTCTGGCATAC | downstream region to construct murAA-gfp |
| murAA_gfp_3pR | TTTCGCTTCCATTCAAAATCGG | downstream region to construct murAA-gfp |
| KanR-R1 | CGCCGTATCTGTGCTCTC | rfp-kanR or gfp-kanR from pACYC-rfp-kan or |
| Rfp_kan-F | GTCGACAGCTA GC CTTATTAACG | rfp-kanR from pACYC- |
| Gfp_kan-F | ACTACTAGTACATAAGGAGGAACTAC | kanR from pACYC-gfp- |

**Table S9.**

List of strains.

| Strain ID | Parent | Genotype | Notes |
|---|---|---|---|
| CAG74209 | *B. subtilis* 168 | *trpC2, lacA::*Pxyl-*dcas9(Erm)* | Peters et al. 2016 |
| CAG78920 | *B. subtilis* 168 | *trpC2, lacA::Pxyl-dcas9(Erm), thrC::*Pveg-*gfp(Spc)* | |
| CAG78921 | *B. subtilis* 168 | *trpC2, lacA::Pxyl-dcas9(Erm), thrC::*Pveg-*rfp(Spc)* | |
| CAG78922 | *B. subtilis* 168 | *trpC2, lacA::Pxyl-dcas9(Erm), sacA::Pveg-rfp* | kan flipped out from sacA::Pveg-rfp(Kan) |
| CAG78923 | *B. subtilis* 168 | *trpC2, lacA::Pxyl-dcas9(Erm), sacA::*Pveg-*rfp, murAA-gfp(Kan)* | |
| CAG78924 | *B. subtilis* 168 | *trpC2, lacA::Pxyl-dcas9(Erm), sacA::*Pveg-*rfp, murAA-gfp(Kan), thrC::*Pveg-*murAA*(Spc)* | |
| CAG78830 | *E. coli* BW25113 | *Tn7att::*P*lLac-O1-dcas9(Gent)* | |
| CAG78108 | *E. coli* BW25113 | *Tn7att*::P*BBa_J23105-dcas9(Gent), yjaA:Pveg-gfp(Cat)*:*yjaB* | |
| CAG78107 | *E. coli* BW25113 | *Tn7att*::P*BBa_J23105-dcas9(Gent), yjaA:Pveg-rfp(Cat)*:*yjaB* | |

**Table S10.**

Table of values for the *murAA-gfp* transcriptional fusion validation experiments.
